# BasePlayer: Versatile Analysis Software for Large-scale Genomic Variant Discovery

**DOI:** 10.1101/126482

**Authors:** Riku Katainen, Iikki Donner, Tatiana Cajuso, Eevi Kaasinen, Kimmo Palin, Veli Mäkinen, Lauri A. Aaltonen, Esa Pitkänen

## Abstract

Next-generation sequencing (NGS) is being routinely applied in life sciences and clinical practice, where the interpretation of the resulting massive data has become a critical challenge. Computational workflows, such as the Broad GATK, have been established to take raw sequencing data and produce processed data for downstream analyses. Consequently, results of these computationally demanding workflows, consisting of e.g. sequence alignment and variant calling, are increasingly being provided for customers by sequencing and bioinformatics facilities. However, downstream variant analysis, whole-genome level in particular, has been lacking a multi-purpose tool, which could take advantage of rapidly growing genomic information and integrate genetic variant, sequence, genomic annotation and regulatory (e.g. ENCODE) data interactively and in a visual fashion. Here we introduce a highly efficient and user-friendly software, BasePlayer (http://baseplayer.fi), for biological discovery in large-scale NGS data. BasePlayer enables tightly integrated comparative variant analysis and visualization of thousands of NGS data samples and millions of variants, with numerous applications in disease, regulatory and population genomics. Although BasePlayer has been designed primarily for whole-genome and exome sequencing data, it is well-suited to various study settings, diseases and organisms by supporting standard and upcoming file formats. BasePlayer transforms an ordinary desktop computer into a large-scale genomic research platform, enabling also a non-technical user to perform complex comparative variant analyses, population frequency filtering and genome level annotations under intuitive, scalable and highly-responsive user interface to facilitate everyday genetic research as well as the search of novel discoveries.

## Introduction

Next-generation sequencing has been widely adopted in life sciences, resulting in massive amounts of DNA and RNA sequencing data available for research and clinical practice (1). The advent of third-generation and single-cell sequencing will contribute both heterogeneity and scale to this already substantial pool of data (2, 3). Consequently, the bottleneck of biological discovery has moved to the interpretation and management of large high-throughput data sets, manifesting especially in the search for common variants in common diseases and pan-cancer genomics, both of these fields often involving thousands of samples and millions of variants per experiment (4, 5). Moreover, the biomedical field is afflicted by a shortage of bioinformaticians with respect to the amount of data analysis and interpretation at hand. Well established workflows, such as Broad GATK, have facilitated primary analysis of the sequence data (6). However, challenges in downstream analysis of variant data, manifesting in non-coding genome and with massive data in particular, have yet to be tackled; the current state-of-the-art in genetics and genomics research and clinical work often involves multiple steps requiring technical expertise for example to implement analysis workflows in scripting languages. For this reason, many individuals may be involved in the analysis of a particular dataset, easily leading into errors due to miscommunication or incompatible skill sets. It is thus desirable to equip the analyst with a comprehensive yet intuitive set of tools to complete the discovery or diagnostics task at hand.

To this end we here introduce a cross-platform graphical software, BasePlayer (http://baseplayer.fi), which is designed to streamline biological discovery by enabling researchers to perform comparative genomic variant studies with human or any other sequenced and annotated reference genomes. BasePlayer is aimed at the analysis of both coding and noncoding regions without expertise in bioinformatics or programming. A brief introduction video of the software can be viewed at (https://youtu.be/kH3QfX868nE). BasePlayer implements an integrative, visualization-driven approach to variant discovery summarized in the following three core features.

First, BasePlayer manages large case-control variant studies in both germline and somatic settings by interactively combining variant comparison, quality and allele frequency filtering, and annotation with genes, pathogenicity predictions, regulatory or user defined regions (Figure 1). These tasks are achieved through a highly scalable and low latency genome browser with multitrack views designed for massive variant data sets of up to thousands of NGS samples (Figure 2 and Supplementary Figure S1). Efficient internal data structures allow high memory and computational efficiency: an ordinary desktop computer with 1 GB of memory allocated to Java Virtual Machine is able to analyze and visualize, for instance, over 3 million somatic variants in over 200 whole-genome sequencing samples. Sequencing, variation and annotation data are presented in space-efficient and resizable overlay tracks.

**Figure 1.**
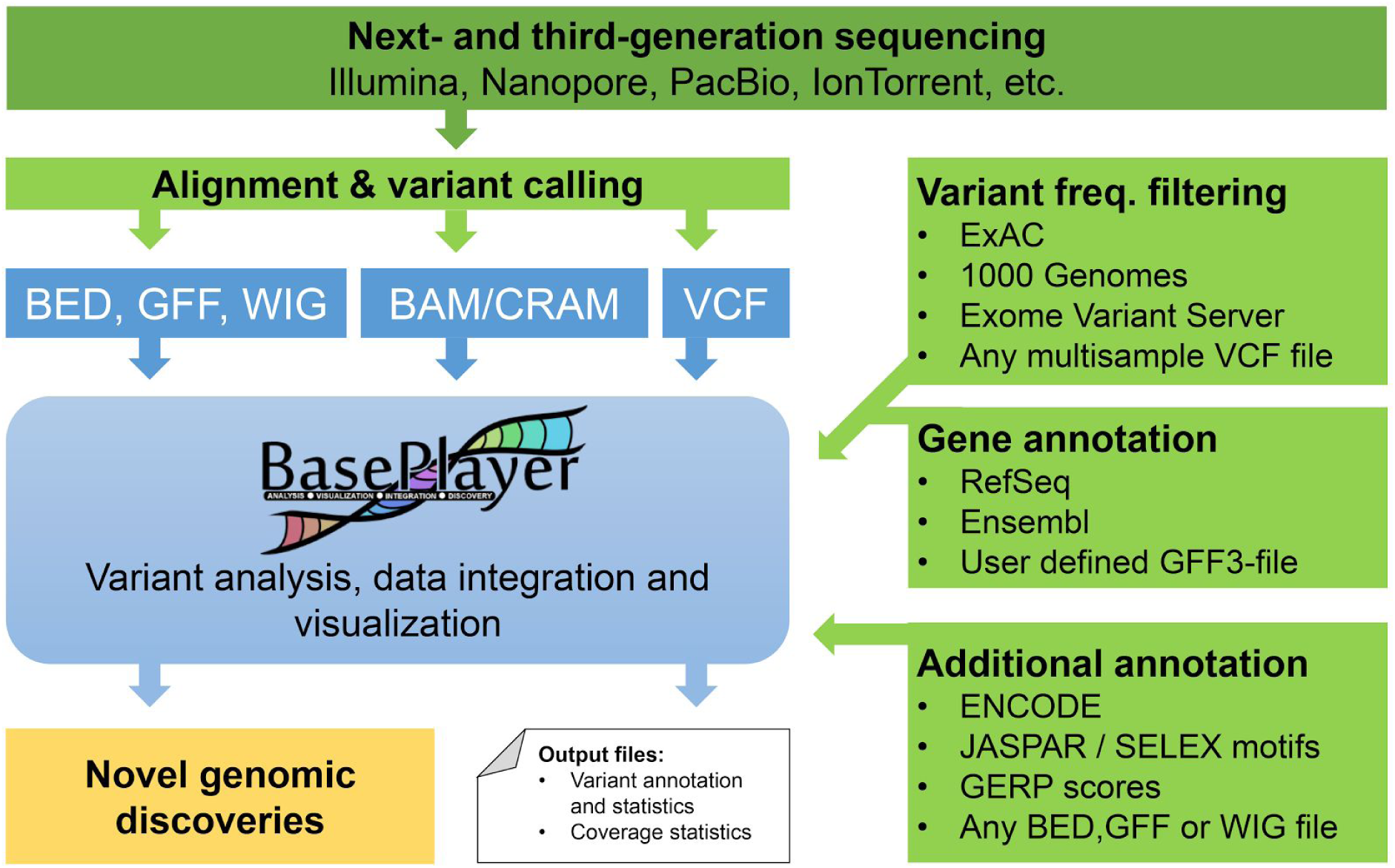
BasePlayer analysis workflow. BasePlayer integrates variant, sequence and annotation data from different sources in standard formats (e.g., VCF, BAM, BED). Preliminary sequence analysis (sequence alignment and variant calling) is performed by the data provider or workflows such as GATK Best Practices prior to BasePlayer analysis to produce these files. Reference sequence and genomic annotations for a given organism can be downloaded from Ensembl or RefSeq (FASTA and GFF3 files). Results may be written in tab-separated (TSV) and VCF formats.

**Figure 2.**
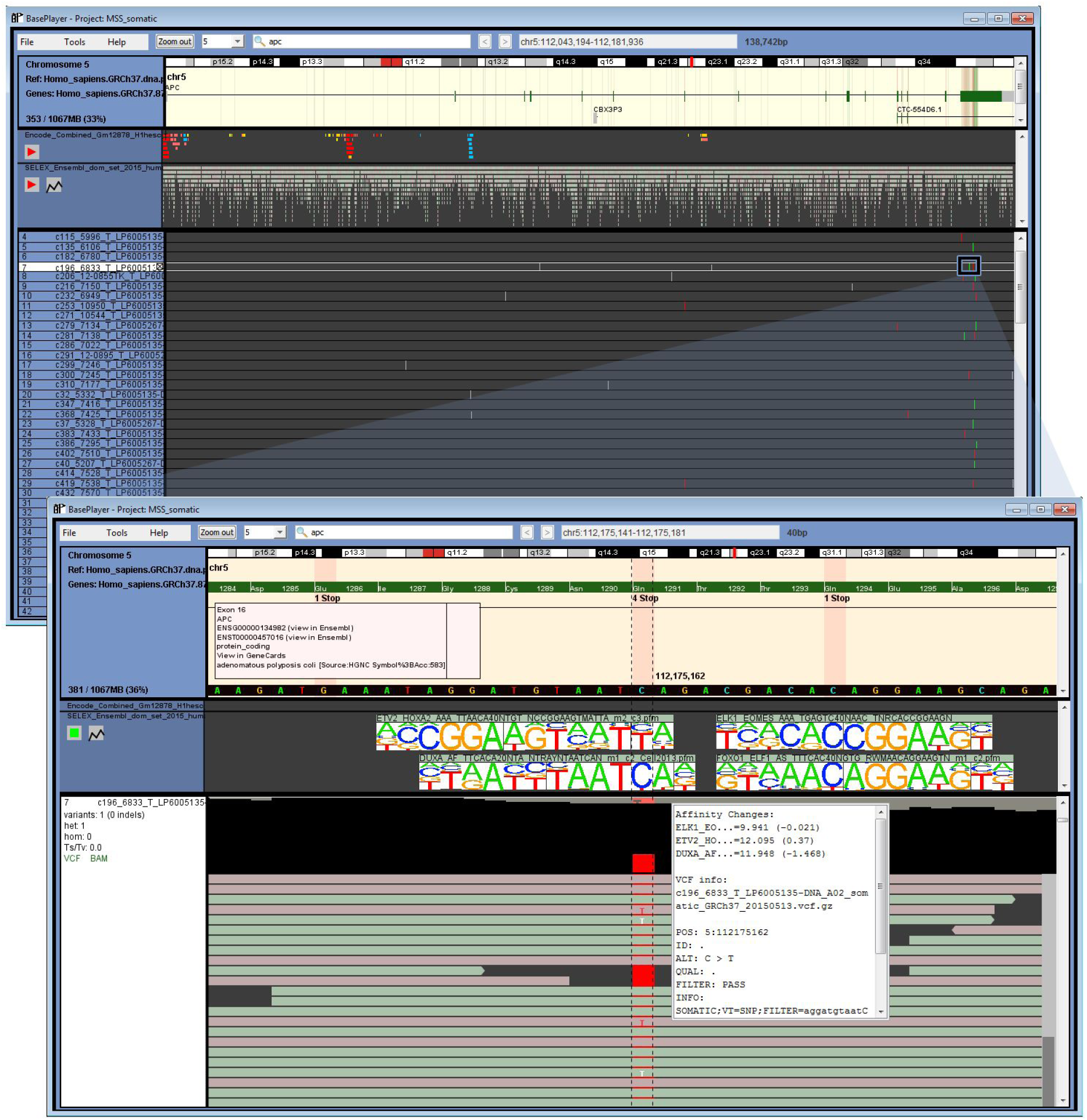
Somatic mutations in *APC* gene, annotation and sequence logo visualization. Upper panel: Gene annotation track visualizes exons and introns of *APC* including collapsed variant view for all opened samples at this locus. Red and green lines indicate hits in exons (SNVs and indels, respectively), grey lines hit in introns, UTRs or intergenic regions. Information box and isoforms can be expanded by clicking the exon. Additional annotation tracks visualize ENCODE regions and TF binding sites and their binding motifs (SELEX) at this locus. Promoters, enhancers and CTCF sites are colored as red, yellow and blue, respectively. Lower panel (zoomed): gene annotation track displays amino acids in exons and possible amino acid changes, derived from base changes in *VCF* files (n=190). In this case, variant hits both in the middle of putative TF binding sites of and in the last exon of *APC*, causing stop-gain. Affinity change values are listed in the variant info box. Sample track visualizes read sequence (BAM) file overlaid with the respective VCF file. Coverage and call information are shown on top of the read and variant panel.

Second, BasePlayer directly supports next- and third-generation sequencing data in BAM and CRAM formats with intuitive visualization of long reads and efficient split views for reads spanning multiple genomic locations (Figure 3 and Supplementary Figure S2) (7, 8). In contrast to genome viewers such as IGV and Savant, variant annotation and comparative analysis at whole-genome level are core functionalities in BasePlayer (9, 10). Furthermore, external server or cloud based analysis platforms, such as BaseSpace (https://basespace.illumina.com/) and Chipster (11), require that the data resides in the cloud, complicating analysis of sensitive or massive data. To address such scenarios, BasePlayer is fully portable, and neither active network connection nor registration is needed. Third, investigation of non-coding variants is facilitated by for instance detection of recurrently or densely mutated genomic regions. By integrating JASPAR, SELEX and variant data, BasePlayer is able to visualize position specific scoring matrices as sequence logos and utilize them to predict affinity changes of mutated transcription factor (TF) binding sites (Figure 2) (12, 13; see methods). In addition, it is straightforward to annotate and visualize non-coding variants with genome-wide data such as genomic event data (e.g. ENCODE), conservation scores and replication timing data to provide an intelligible view on the regulatory genome.

**Figure 3.**
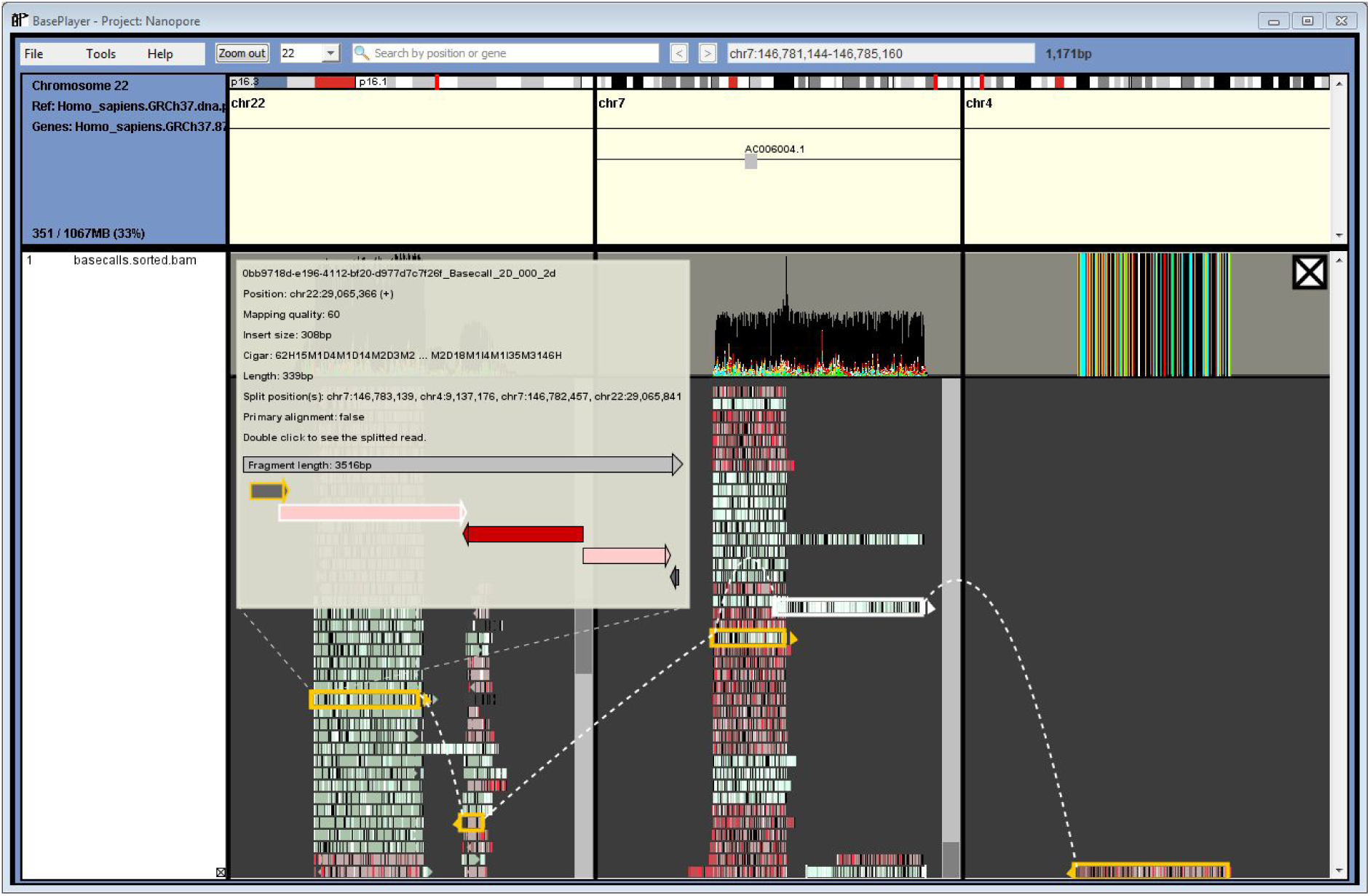
BasePlayer visualization of third-generation sequencing data. A single read of 3,516 bp and its component mappings have been highlighted. The read sequence has been mapped to three chromosomes (22, 7 and 4), which is visualized with multiple split views. Also, the relative position and orientation of the mappings are visualized in the info box.

## Results

### The search for pathogenic inherited variant with exome sequencing data

We demonstrate the analysis capabilities of BasePlayer with three different examples, each performed with an ordinary desktop computer and with 1 GB of allocated memory. First, we show a workflow for finding the predisposing mutation in a family with an inherited disease (meningioma), replicating the previous study described in (15). The aim of this study was to find a causative genetic variant able to explain the clustering of meningioma cases in a family of five affected siblings (15). The data set included three exome sequenced germline samples from the affected siblings as well as genome-wide linkage analysis data of the same family.

We set out to find the causative, non-synonymous mutation which is assumed to segregate in the family, reside in the linkage-compatible region and have a very low allele frequency in the general population. This study demonstrates the utility of sample comparison features, gene and linkage annotation, and the use of publicly available control data sets (here we use data from the ExAC; see methods) (17). Real-time demonstration video for this case can be viewed at https://youtu.be/zCUBY-IdY2I. Required files for this study are variant files (VCF) for cases and controls, BAM files for read level inspection and a BED file for the linkage-compatible regions. First, the VCF files for all three samples are opened in separate tracks (Figure 4A), and minimum variant quality and coverage thresholds are set by the user. BasePlayer visualizes BAM and VCF files of the same individual/case within the same track, in contrast to many other tools. Second, the linkage-compatible regions are opened as a new track, which is applied to remove variants in the genomic regions that are not segregating in the affected individuals (Figure 4B). Variants which are not shared by all individuals are then excluded using the ‘Common variants’ slider (Figure 4B, top right corner). Next, ExAC variants, excluding TCGA data (n=8,608,691 entries in 53,105 individuals; multisample VCF file) are opened as a control track. The allele frequency threshold is set to 0.001 and when applied, all variants that are present more frequently in the control data are filtered (Figure 4C). Finally, BasePlayer calculates non-synonymous variants for every chromosome with all used filters and settings applied. The result contains a total of 9 variants in 9 genes with these settings. Of these, a missense variant in *SUFU* is the only one that does not appear in the ExAC data and is a prime candidate for the causative gene, a result which has been replicated in later studies (15). BasePlayer is thus able to complete familial candidate gene discovery in a few minutes from VCF files without use of scripting or command-line tools.

**Figure 4.**
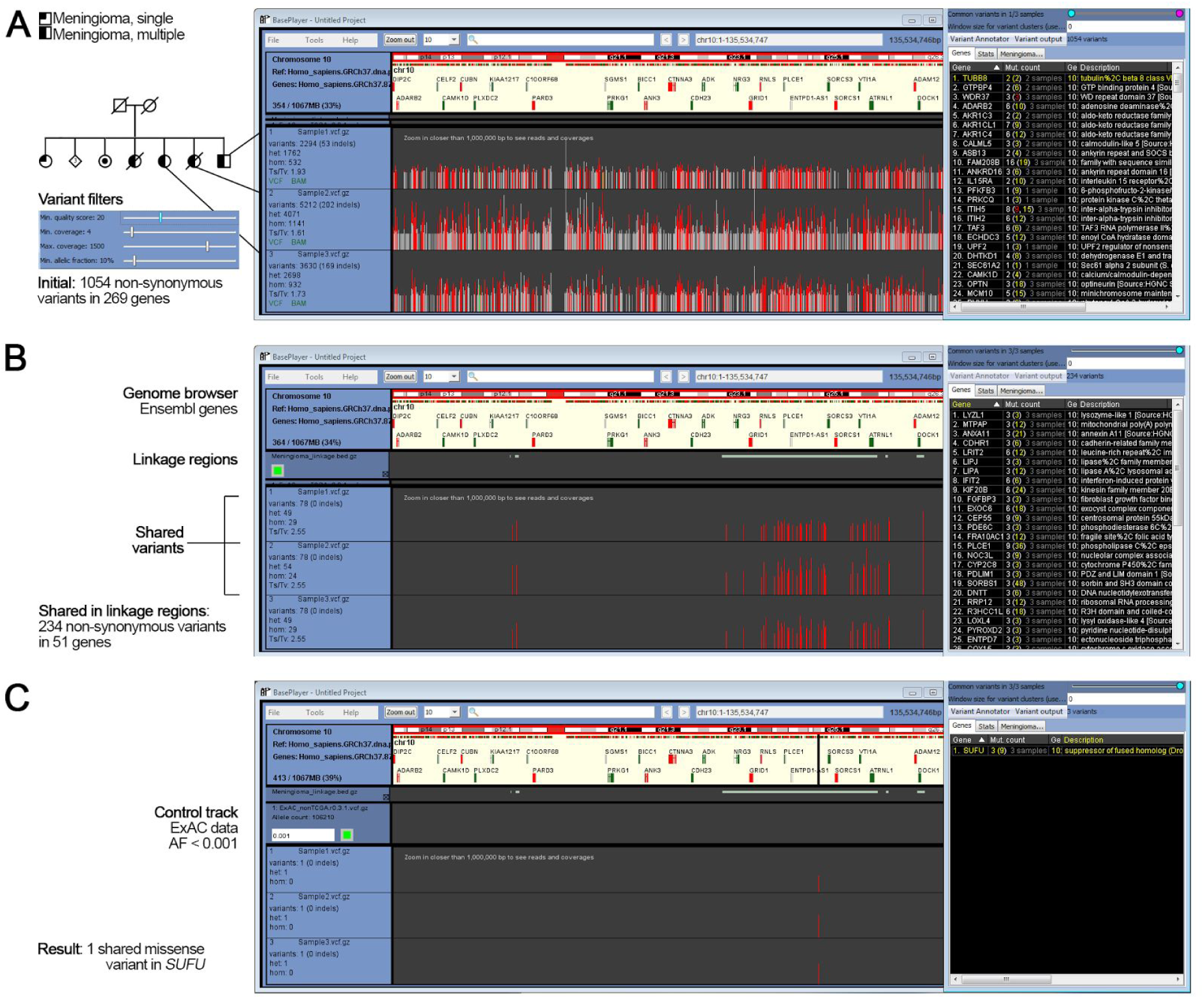
Familial study workflow with BasePlayer. (**A**) Three exome sequenced germline samples and their variants in chromosome 10 visualized in the main window of BasePlayer. Vertical lines represent variant calls in a sample, colored as red, green (SNV or indel in coding region, respectively) and grey (variant in non-coding region). Height is relative to the coverage at variant locus. Variant filters (left panel) are used to exclude low quality variants. The right panel shows a list of all genes in chromosome 10 with non-synonymous mutations in this data set. (**B**) Linkage regions are shown in a separate track below genome browser. Non-coding variants have been filtered out from sample tracks. ‘Common variants’ slider in the top-right corner has been used to compare variants between samples. Only variants shared between all samples (3/3) have been retained. (**C**) ExAC variant filtering has been applied to exclude common polymorphisms.

### Somatic mutation clusters in regulatory genome

Second case describes variant analysis in noncoding cancer genomes. Here we describe an approach to find mutations with a possible role in tumor development by detecting somatic mutation clusters in annotated regulatory regions and TF binding sites. The study is performed with somatic variant calls of 190 whole-genome sequenced colorectal cancers (16). ENCODE regulatory data is used as an annotation track (14; see methods). Real-time demonstration video for this analysis can be viewed at https://youtu.be/D2c-57EPYJg. First, VCF files are opened and BasePlayer displays variants for the selected chromosome (Figure 5A). Variant filter thresholds are set to minimum to include low quality calls, resulting in a total of 3,012,306 variants across all chromosomes. Variant clustering is performed by setting the window size to 100 bp and the minimum number of variants in a cluster to four (Figure 5B). Variants are then intersected with the regulatory regions. With these settings, genome-wide annotation took less than two minutes to process and resulted in 665 variant clusters. Highest variant count (20) was observed at CTCF region in chromosome 20 in the intron of *CBFA2T2* gene (Figure 5B). The prediction of possible pathogenicity of non-coding mutations can be facilitated by utilizing e.g. CADD, GERP and transcription factor binding specificity data. Analyzing sequence conservation and motifs alongside the mutation data can reveal sites, where mutations tend to change predicted affinities of TF binding more frequently than expected (Figure 2 and 5C). The knowledge from this “second genetic code”, containing predicted TF binding sites, provides a valuable asset to unravel mechanisms behind the regulation of gene expression.

**Figure 5.**
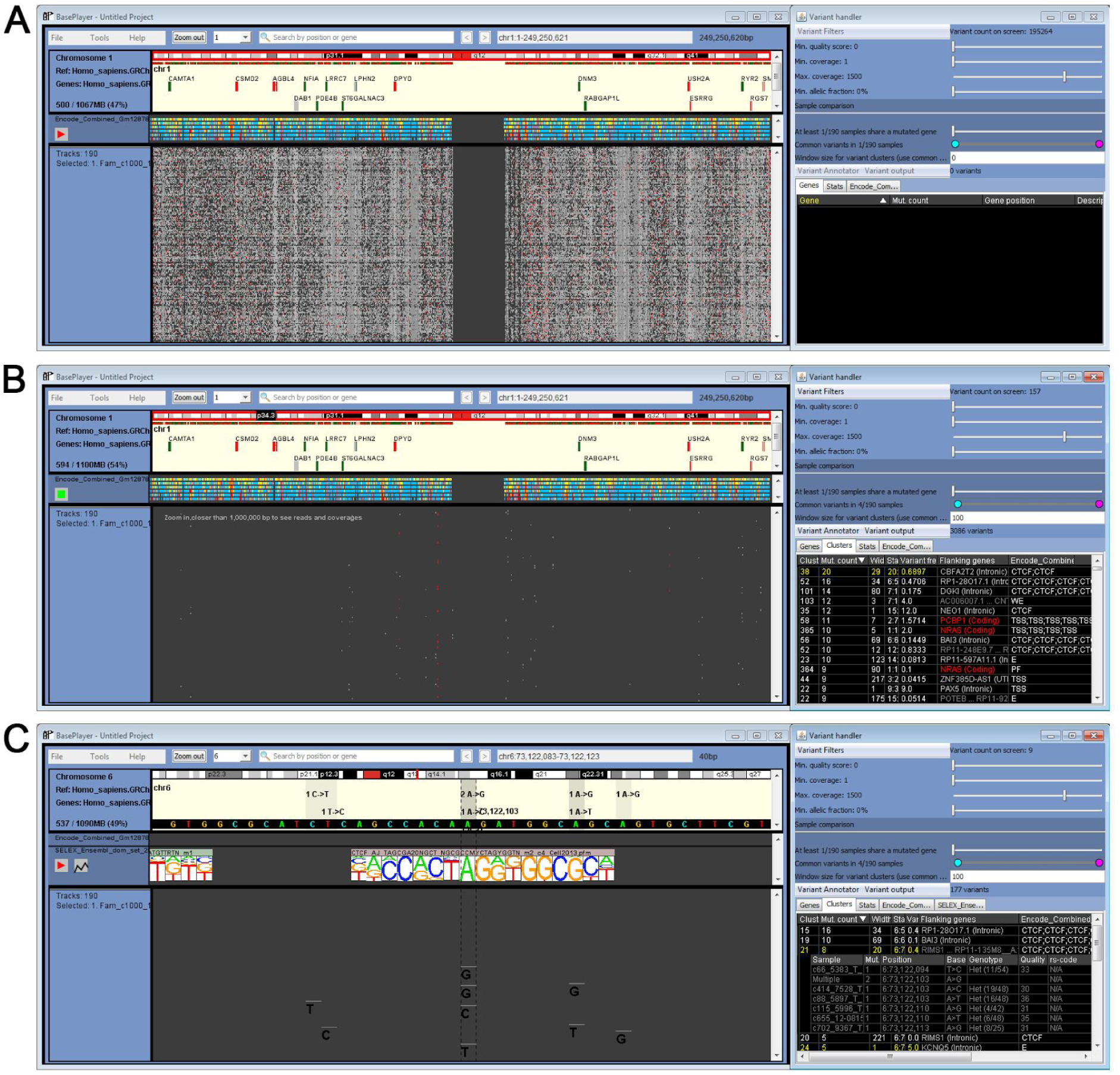
Somatic variant clusters in regulatory genome. (**A**) All somatic single nucleotide variants of 190 colorectal cancers (WGS) in chromosome 1 are visualized. Additional track in the middle visualizes ENCODE regulatory regions, colored as blue: CTCF binding sites, dark yellow: enhancers, light yellow: weak enhancer, red: transcription start site, light red: promoter flank. All variant filtering thresholds have been set to minimum (right panel). (**B**) ENCODE track has been activated, which annotates and excludes all non-overlapping variants from the study. On right panel, all clusters overlapping regulatory regions have been calculated and sorted by the variant count. (**C**) Motif visualization with JASPAR and SELEX data. Closer inspection of the cluster highlighted in (16) shows mutation hotspots at an occurrence of CTCF binding motif.

### Detection of retrotransposon target sites with long-read data

Third, the advent of third-generation sequencing (e.g., Oxford Nanopore, Pacific Biosciences) has enabled studies of genomic regions and events that were not feasible to analyze with short-read data. Thousands of base pairs long reads require flexibility from the visualization software, as single read can span multiple genomic breakpoints, or multiple exons in the case of RNA data. In BasePlayer, the researcher can divide the main screen between distinct genomic loci by double clicking a read that has been mapped to multiple locations (Figure 3). Demonstration video for visualizing long-read data in BasePlayer can be viewed at https://youtu.be/E7SSKxzn4vk. All split views are equally functional and contain gene annotation (e.g., in search of fusion or target genes). In addition, BasePlayer visualizes whole DNA fragment and relations of its splitted parts in an inset info box (Figure 3).

## Discussion

Large-scale data integration of burgeoning sequence, variation and functional data has never been as relevant as it is today. Regulatory genomics, for example, often require joint analysis of genomic data from different experiments such as ChIP-seq/exo/nexus, RNA-seq, ATAC-seq and ChIA-PET (19). Added value lies in the integration and comparison of various data types, where unexpected or already familiar phenomena can emerge in a new light (Supplementary Figure S3). Thus, the value of genome-wide variant data increases together with accumulating functional knowledge. BasePlayer provides means to apply this knowledge in practice to drive biological discoveries - its previous, unpublished version (RikuRator) has already been successfully used in several efforts (Supplementary Table S1). While most of large-scale comparative variant analysis and data integration occurs in server environments using various tools and scripting languages, BasePlayer contributes to a paradigm shift of genomic variant discovery to more interactive, practical and visual direction. By doing so it also represents a breakthrough in genome interpretation software that require few specialized skills, paving the way towards tools that can be exploited by the general population.

## Materials and Methods

### Annotation data

Human reference genome (GRCh37) and gene annotation data for the example cases were downloaded from Ensembl database (Release 87). ENCODE data for regulatory regions was published in (14). We merged data from six different cell lines (Gm12878, H1hesc, Helas3, Hepg2, Huvec and K562) to ENCODE annotation file used in Case 2.

Position specific matrices for transcription factors were obtained from (13) and the JASPAR database (12). TF binding sites for GRCh37, obtained from Ensembl Biomart, and ENCODE annotation file can be downloaded from BasePlayer website (downloads).

### Sequence and variant data

Exome-sequence data and variant calling for meningioma patients (Case 1) were described in (15). Whole-genome sequencing data of colorectal cancer samples and identification of somatic variants (Case 2) were described in (16). Third-generation sequencing data (Case 3) was produced with Oxford Nanopore Technologies MinION (SQK-MAP005) and alignment was performed with BWA-MEM 0.7.12-r1039 (7).

ExAC (Exome Aggregation Consortium) variant frequency data (0.3.1) was downloaded via ExAC FTP server (17).

### Variant annotations

Amino acid and gene annotations are calculated by using indexed reference sequence (FASTA), gene annotation (GFF3) and variant (VCF) files. Gene annotation file contains, among others, chromosomal positions and codon phases for all protein coding exons, which are used to fetch codon sequence triplet from reference sequence file. Amino acid change is derived from variant position and base change (reported in VCF file) at fetched codon sequence. BasePlayer annotates synonymous, non-synonymous, nonsense, splice-site, frameshift, inframe, UTR, intronic and intergenic variants, compatible with Annovar annotation (18). Closest flanking genes are reported for intergenic variants.

### Variant clustering

Calculation of variant clusters requires two values set by the user: maximum distance between adjacent variants (window size) and minimum number of variants in the cluster. A cluster is defined as a set of variants, where the distance between any two adjacent variants does not exceed the window size. Thus, the cluster width (distance between the leftmost and rightmost variant) can exceed the window size.

### Additional Java packages

BasePlayer utilizes Htsjdk (version 1.141, https://github.com/samtools/htsjdk) index readers for BAM, CRAM and Tabix indexed files. CRAM reader was optimized for BasePlayer by modifying CRAMFileReader and related classes from the Htsjdk package. Index readers for BigBed and BigWig files were downloaded from https://github.com/lindenb/bigwig,originally written by Martin Decautis and Jim Robinson for the Integrative Genomics Viewer (Broad Institute).

### Software

BasePlayer is able to analyze NGS data in standard formats, including targeted (e.g. exome), whole-genome and RNA sequencing data (Supplementary Figure S4) generated by Illumina, Ion Torrent, Oxford Nanopore and PacBio platforms. Currently supported data formats include indexed VCF, BAM, CRAM, BED, TSV, GFF3 and Wiggle/bigWig/BigBed. RNA-sequencing data is displayed similarly as short-read DNA sequencing data. Reads overlapping an exon boundary are split and visualized accordingly.

For interactive large-scale variant analysis and visualization at least 2 GB of allocated memory for Java VM is recommended. However, BasePlayer is also able to perform variant analysis in batches of genomic windows (default window size 1 Mbp) if necessary with only 1 GB of memory.

## Availability

BasePlayer is available free of charge at http://baseplayer.fi. BasePlayer can be run on Windows, Linux and Mac computers with Java JRE 1.7 or newer. Video tutorials and demonstrations can be viewed at BasePlayer channel on Youtube: https://www.youtube.com/channel/UCywq-T7W0YPzACyB4LT7Q3g. Source code is available at BasePlayer project page on GitHub:https://github.com/rkataine/BasePlayer.

## Acknowledgements

We thank Teemu Kivioja for guidance with SELEX data and Alison Louise Ollikainen for the voice work in demonstration videos. We thank Barun Pradhan and Liisa Kauppi for sharing their unpublished Nanopore data. We also like to thank Mervi Aavikko, Linda van den Berg, Outi Kilpivaara, Johanna Kondelin, Yilong Li, Miika Mehine, Heikki Metsola, Lauri Sipilä, Tomas Tanskanen, Pia Vahteristo and Niko Välimäki for testing BasePlayer and giving suggestions and additional support. Finally, we acknowledge ZeroTurnaround for creating the JRebel plugin for Eclipse (IDE).

